# Sequence effects on mutation rates investigated in whole-genome sequenced UK Biobank participants

**DOI:** 10.64898/2026.03.06.710079

**Authors:** David Curtis

## Abstract

UK Biobank has released whole genome sequence data for 500,000 participants, including allele counts for hundreds of millions of variants and these were considered in the context of the pentanucleotide background on which they occurred. Variants with an allele count of 25 were found to closely mirror previously reported de novo mutations (DNMs) in terms of the frequencies of variant types. Therefore these variants, referred to as AC25 variants, were used to investigate factors relevant to mutation frequency. The counts of AC25 variant types in a trinucleotide context could be well approximated by combining seven mutational signatures previously obtained from studies of cancer cells. Frequencies of variants were strongly influenced by context. C>T variants occurred more often in the CpG context but other features of the trinucleotide context also had marked effects. Although the trinucleotide context was a very strong predictor of the variant frequencies for the full pentanucleotide context, there were examples where the more distant nucleotides did have marked effects. Here, the configuration of bases two upstream or downstream could have a more than two-fold effect on variant frequencies, presumed to reflect mutation rates. For some variants frequencies varied between non-transcribed and transcribed regions and variants with higher frequencies in transcripts of protein coding genes also demonstrated strand asymmetry, being more frequent on the coding than template strand. Genes with high frequencies of these variants showed enrichment for basic cellular processes, meaning that they could have arisen as a result of transcriptional events during embryogenesis prior to entering the germline. Investigating the molecular mechanisms whereby contexts moderate mutation rates could lead to a better understanding of variant pathogenicity and how DNA fidelity is conserved in normal tissue.

This research has been conducted using the UK Biobank Resource.

**Article summary:** We analysed hundreds of millions of variations in DNA code observed in 500,000 UK Biobank participants. For each change of a single DNA base, we also considered the two bases either side. We found that the set of variants which each occurred exactly 25 times in the sample, termed AC25 variants, had a frequency distribution which almost exactly matched that of a set of previously reported variants which occurred as new mutations. Therefore, we used these AC25 variants to study the effect of DNA context on mutation rates and to compare mutation types with those previously observed in cancer patients.

## Introduction

Ultimately, all genome sequences are derived from processes which involve mutation, in which there are imperfections in the molecular processes responsible for copying the genetic code from one cell to another. Investigating patterns of mutation may yield insights into these underlying mechanisms. Additionally, it could assist in distinguishing individual variants which are of rare types and hence which might be more likely to be pathogenic (Curtis, 2024). In cancers, mutations occur with increasing frequency as the disease progresses and a fruitful area of research has been to study the mutational signatures seen in different types of cancer (Alexandrov et al., 2020). This involves enumerating the counts for each mutation, for example a single base substitution, specific to the background upon which that mutation occurs, for example the two flanking bases. The mutations observed in a sample can be considered to be produced by number of different drivers of mutation involved in DNA damage, repair and/or replication, possibly involving different endogenous and exogenous exposures, and each driver will produce a characteristic mutational signature. Methods have been devised to disaggregate an observed set of mutations into its component signatures (Díaz-Gay et al., 2023).

Although cancers provide a useful source of data involving large numbers of somatic mutations, information can also be derived from other sources. If parents and children are both sequenced then de novo mutations (DNMs) can be observed (Jónsson et al., 2017; McRae et al., 2017; Rahbari et al., 2016). A sample of 2,976 whole genome-sequenced trios was used to identify 200,435 DNMs and these were used to assess the mutation probabilities for different trinucleotides as well as how these were impacted by crossover events (Halldorsson et al., 2019).

Whole genome sequenced population samples such as 1000 Genomes can be used to reconstruct genealogies and these can then be used to make inferences about mutational processes and their association with different population features (Gao et al., 2023). Sequencing sperm allows for the investigation of mutation within an individual and can demonstrate the relationship between mutation rate and paternal age (Shoag et al., 2025). An alternative approach is to study polymorphisms observed in a large sample of individuals and in this context it has been reported, for example, that C>T substitutions are seen more often in the context of CpG dinucleotides, an observation which has been explained as being due to the spontaneous deamination of 5-methylcytosine (Aggarwala and Voight, 2016; Zhao and Boerwinkle, 2002). Effects on substitution rates were also observed for more distant bases, with the heptanucleotide context explaining >81% of variability in substitution probabilities (Aggarwala and Voight, 2016). Frequencies of all polymorphisms observed in a population reflect not only ongoing mutational processes but also historic processes, including selection. Instead, attention may be confined to variants occurring only as singletons, on the assumption that these are likely to reflect patterns of recent mutation. In a sample of 3,560 whole-genome sequences for the Bipolar Research in Deep Genome and Epigenome Sequencing (BRIDGES) study, 36 million singleton variants were used to estimate relative mutation rates for different mutation types, which were shown to be influenced by nucleotide context and other genomic features (Carlson et al., 2018).

More recent work has thrown into question the appropriateness of using singleton variants to estimate mutation rates. With a constant population size and a neutral mutation rate *u*, the allele frequency spectrum is such that the expected number of variants to have exactly *i* copies of the alternate allele in a sample is proportional to *<u>u</u> / i* under the “infinite sites” assumption that there are no recurrent mutations (Kimura, 1969). However for sites with high mutation rates at which recurrent mutations can occur there will be a reduction in low frequency variants and an excess at higher frequencies (Wakeley et al., 2023). This means that the singleton count is low relative the mutation rate for variant types whose mutation rate is high and so overall the singleton counts across different variant types will not correlate well with their mutation rates. A follow-up study of the BRIDGES samples confirmed that the allele frequency spectrum differed between variant types, with the singleton to doubleton ratio being lower for variant types with higher mutation rates, and showed that this heterogeneity could impact inferences about population genetic history (Liao et al., 2023).

Although mutations can occur through errors in DNA replication or failures of repair mechanisms following damage, transcription associated mutations can be considered separately and these can lead to strand asymmetries (Cui et al., 2012; Green et al., 2003). In order for mutations produced during transcription to be observed in the germline, they must occur in genes which are expressed early in embryonic development or in the lineage of gametogenic cells. Whole genome sequence data for 500,000 UK Biobank participants has now been released, including hundreds of millions of variants (The UK Biobank Whole-Genome Sequencing Consortium., 2025). This large sample was used to obtain estimates of relative mutation rates for different types of variant according to their pentanucleotide context. Mutation rates were compared between transcribed and non-transcribed regions and for transcribed regions additional analyses were carried out to identify mutation types demonstrating strand asymmetry.

## Materials and Methods

### Obtaining variant counts

The UK Biobank Research Analysis Platform was used to access the whole genome sequence data which had been obtained using the methods previously described (The UK Biobank Whole-Genome Sequencing Consortium., 2025). UK Biobank had obtained ethics approval from the North West Multi-centre Research Ethics Committee which covers the UK (approval number: 11/NW/0382) and had obtained written informed consent from all participants. The UK Biobank approved an application for use of the data (ID 51119) and ethics approval for the analyses was obtained from the UCL Research Ethics Committee (11527/001). For each chromosome, the pvar file was downloaded for the DRAGEN population WGS variants in PLINK format. The pvar file contains the coordinate, DNA change and allele count for each variant but no information about individual participants.

Attention was restricted to single base substitution (SBS) variants. All analyses were confined to non-centromeric regions because the centromeres include repetitive DNA sequences which are highly homogeneous and hence their sequence content is not representative of the rest of the genome (Altemose et al., 2022). The telomeric regions were also excluded. In the included regions there were a total of 1.06e9 SBS variants, of which 4.05e6 were singletons and 1.48e6 were doubletons.

Each SBS variant was considered in the context of the two bases 5’ and two bases 3’, so that the fundamental unit of analysis consisted of a pentanucleotide sequence with the variant consisting of a change from reference to alternate for the central base. A variant type is then defined by six bases, the pentanucleotide background along with the alternate central base, for example AA[G>C]TG, meaning that there was a total of 4^5*3 = 3,072 types. The allele count (AC) field for each pvar file was used to obtain total counts for each variant type binned according to the AC field, consisting of counts of singletons, doubletons, AC = 3, etc. up to AC=200.

A list of DNMs obtained from 2,786 whole genome sequenced trios described above was downloaded from https://www.science.org/doi/suppl/10.1126/science.aau1043/suppl_file/aau1043_datas5_revision1.tsv (Halldorsson et al., 2019). Counts of variant types among these DNMs, consisting of the variant in the context of its pentanucleotide background, were enumerated. In the regions excluding the centromeres and telomeres, there were a total of 178,610 SBS DNMs.

For each AC bin, the frequency of each variant type was obtained, consisting of the count divided by the number of times the background pentanucleotide occurred in the reference genome, and the frequencies for the DNMs were obtained in the same way. As previously reported, the ratio of singletons to doubletons was markedly reduced for variant types with higher mutation rates, consisting of those including the [C>T]G motif, and across the 3,072 types the correlation of singleton frequencies with the DNM frequencies was only 0.043. This correlation increased to 0.442 for doubletons and to 0.704 for tripletons and continued increasing with allele count. At the same time, the total number of variants in each AC bin gradually reduced. With AC=25, the correlation between variant frequencies and DNM frequencies across variant types reached 0.990. Thus, the variants which have an allele count of 25 almost exactly predict the relative mutation frequencies observed directly in DNMs and these will now be referred to as AC25 variants. Classical population genetics predicts that the expected number of variants to have *i* copies of the alternate allele is proportional to *<u>u</u> / i* so that for a particular value of *i* the relative frequencies of variant types are expected to mirror their mutation rates (Kimura, 1969). While this breaks down for singletons and other rare variants, presumably because of recurrent mutations, as the allele count increases towards 25 it seems that the distortion becomes negligible and the frequencies become accurate indices of relative mutation rates. There were in total 2,654,130 AC25 variants so it was thought that this sample, much larger than the 178,610 DNMs, could be used to obtain more accurate estimates of effects of sequence on mutation probability than had been possible with the DNMs.

### Mutational signature analysis

Previous investigations of mutational signatures in cancer have used counts of variants occurring on a trinucleotide background sequence and use the 96 variant types having C or T as the central base, using the sequence on the minus strand when the central base on the plus strand is G or A (Alexandrov et al., 2020). These counts depend on both the mutation rate and the number times the background sequence occurs within the genome. Hence, the AC25 counts for the variant types based on pentanucleotide backgrounds were used and were pooled according to this scheme using the central trinucleotide background. Sets of mutation-type specific reference signatures have been developed and deposited in the Catalogue of Somatic Mutations in Cancer (COSMIC) database(Alexandrov et al., 2020; Sondka et al., 2023; Tate et al., 2019). These signatures were downloaded from the COSMIC website at https://cancer.sanger.ac.uk/signatures/downloads/. Code was written to combine these signatures linearly to best fit the observed AC25 counts for the 96 types, with coefficients constrained to be positive (by fitting the square roots of the coefficients). Those signatures for which the estimate of beta/SE exceeded 7 were retained in the model which was then refitted, to produce a parsimonious model to predict the observed AC25 counts from a small number of reference signatures.

### Modelling effects of background on variant frequencies

In order to allow for separate analyses of patterns of mutation related to transcription, the genome was divided into transcribed and non-transcribed regions. For the non-transcribed region variants were specified according to the plus strand whereas for transcribed regions variants were specified according to coding strand. The transcribed region was further subdivided between transcripts for protein coding genes and transcripts for non-coding RNA genes, since the coding genes are transcribed by RNA polymerase II while RNA genes are also synthesized by RNA polymerase I and III (Bunch, 2018). There were 1,457,448 AC25 variants in the non-transcribed region and their frequencies correlated at R = 0.99 with the frequencies which had been observed in the DNM sample (Halldorsson et al., 2019). The frequencies also correlated at R = 0.99 with those which had been obtained for European subjects in the 1000 Genomes Project and which had been used for the earlier study of the effect of sequence context on substitution probabilities (Aggarwala and Voight, 2016). These AC25 variants in the non-transcribed region were taken to result from mutations not due to transcription and were used for the analyses of effects of sequence context on mutation probabilities.

The general approach to analysing sequence effects on mutation probabilities involved studying the frequency of observing a certain alternate allele conditional on the reference sequence background. For example, if the T nucleotide occurs 7.78e8 times in the reference genome and we observe 3.23e5 AC25 T>C variants then we say that the frequency of AC25 T>C variants occurring on the background of the T nucleotide is 3.23e5/7.78e8 = 0.000415. Likewise, if the trinucleotide sequence ATG occurs 4.55e7 times in the reference and there are 4.00e4 AC25 A[T>C]G variants then the frequency of AC25 A[T>C]G variants on the ATG background is 4.00e4 / 4.55e7 = 0.000880. Finally, if the pentanucleotide CATGC sequence occurs 3.89e6 times in the reference and there are 2,680 CA[T>C]GC AC25 variants then the frequency of CA[T>C]GC AC25 variants on the CATGC background is 2,680 / 3.89e6 = 0.000689.

Using this approach, it is possible to implement a logistic regression framework whereby the outcome to be predicted is a variant with a specific alternate base, referred to as the target, and this outcome is modelled as being influenced by the background reference sequence. Where background sequences are mutually exclusive, for example when considering all trinucleotides consisting of a central base and the flanking base on each side, one can simply total up counts across matching pentanucleotides to obtain the frequency for each background to produce a AC25 variant yielding a particular target base. This process was carried out for single nucleotides, dinucleotides including one upstream base, trinucleotides including two upstream or two flanking bases and tetranucleotides including two upstream and one downstream base.

In order to explore the effect on mutational probabilities of including the two flanking nucleotides relative to the effect of the core nucleotide, AC25 variant frequencies were obtained for background core nucleotides and then each flanking trinucleotide sequence was included in turn in a logistic regression model to estimate its additional effect on the odds of producing the target base. Similar analyses were done to measure the effects of models with the full pentanucleotide sequences relative to the flanking trinucleotide sequences.

### Analysing effects related to transcription

The frequencies of AC25 variants in coding transcripts and non-coding transcripts both correlated with the frequencies for variants in non-transcribed regions with R > 0.995. However there were some variants for which frequencies were slightly but statistically significantly different. To investigate these differences, a logistic regression analysis was implemented to model whether the frequency of a variant differed between one region and another. Three comparisons were performed: between the non-transcribed region and protein coding gene region, between the non-transcribed region and RNA gene region and between the protein coding gene region and the RNA gene region. Analogously to the modelling of mutation probabilities, for each comparison modelling was performed using effects of single nucleotides and then incorporating varying numbers of flanking nucleotides up to the full pentanucleotide sequence.

A similar approach was used to model effects on strand asymmetries, with the outcome of interest being whether the variant occurred on the plus or minus strand for the non-transcribed region or whether it occurred on the coding or template strand for the transcribed regions.

In order to assess which genes were most affected by transcription related mutational processes, variants were identified which were more frequent in transcripts of protein coding genes than the non-transcribed region and which showed greater strand asymmetry in that they had higher frequency on the coding strand than the template strand. For these gene-based analyses, all variants were used rather than only those with an allele count of 25. The 500 genes with the highest frequencies of these variants were entered into the PAN-GO gene ontology enrichment analysis to determine which biological processes they were involved with (https://functionome.geneontology.org/) (Mi et al., 2013).

### Data analysis

Code to count variants and background sequences as well as to carry out modelling was written in C++ and is available at: https://github.com/davenomiddlenamecurtis/countDNAVariants. This repository also contains raw counts and calculated frequencies of the variants analysed. Additional data management and figure preparation was performed using R v4.3.2 (R Core Team, 2014), including the scico package to utilise Fabio Crameri’s scientific colour maps (DOI 10.5281/zenodo.1243862).

## Results

### Mutational signature analysis

For the mutation signature analysis, the counts of standard 96 mutation types, consisting of all trinucleotide AC25 variants in which the core base was either C or T, were considered with the counts from the plus and minus strands being combined. As described in the Methods section, 86 reference signatures obtained from the COSMIC database were entered into a multiple linear regression analysis to obtain coefficients for a linear combination of signatures which would provide the best fit to the observed counts of AC25 variants.(Tate et al., 2019). Coefficients were constrained to be positive and the intercept was constrained to zero. When this was done there were seven signatures for which the ratio of the coefficient to its standard error exceeded 7 and these signatures were retained and the modelling repeated. These seven signatures along with their best fitting relative contributions are shown in Table 1. It should be noted that the proposed aetiologies for these signatures as described in the COSMIC database do not necessarily reflect the mutational mechanisms which have produced them (Sondka et al., 2023). In particular, although the SBS87 signature can result from thiopurine chemotherapy treatment, in the present context we would interpret this as saying that SBS87 might result from similar biological mechanisms as those produced by thiopurine rather than resulting from actual exposure to thiopurine. The observed counts of mutation types and the predicted counts which would be obtained from combining these signatures according to the best-fitting weights have correlation coefficient R = 0.993. These observed and predicted counts are shown in Figure 1.

**Figure 1.**
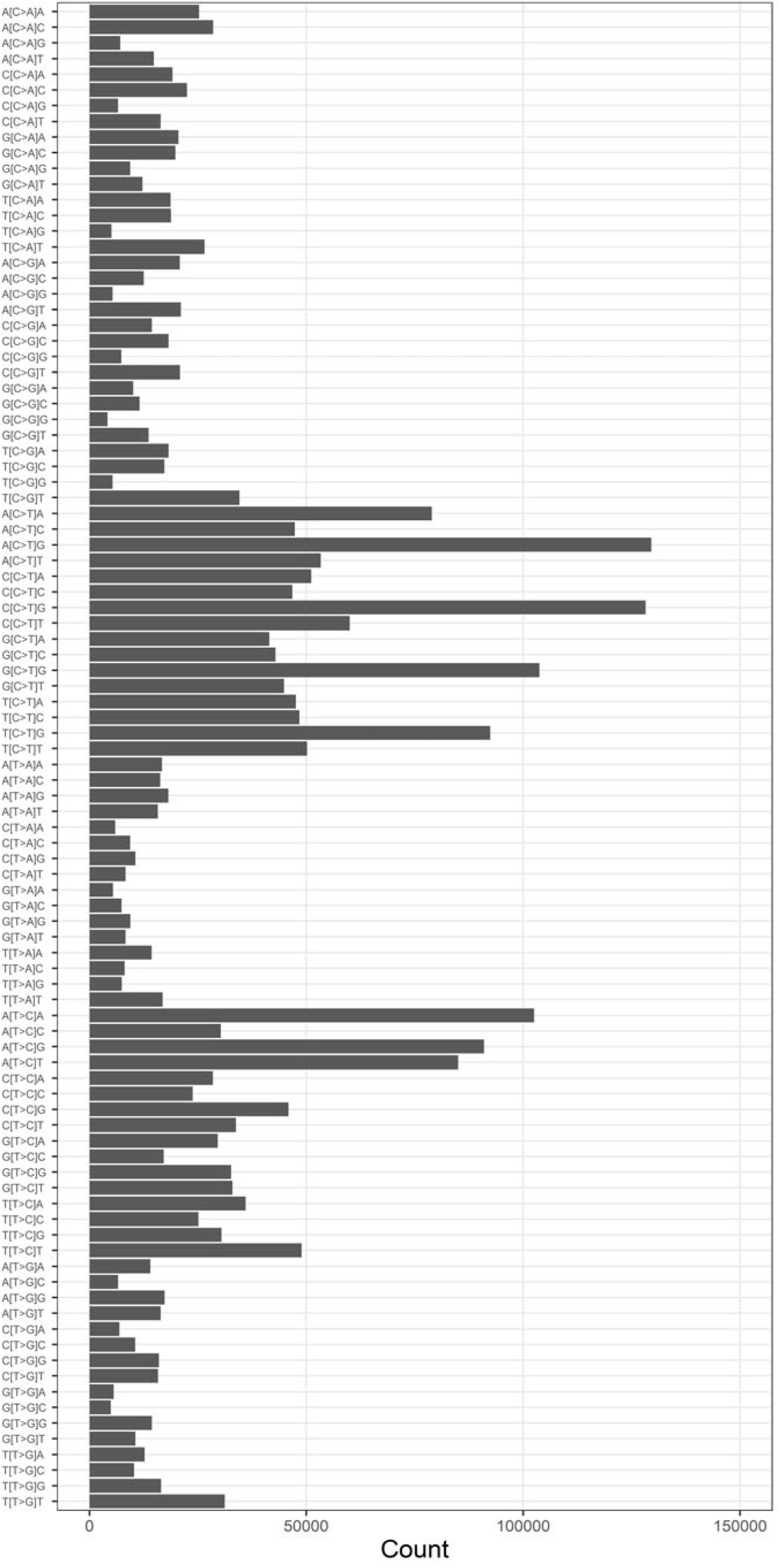
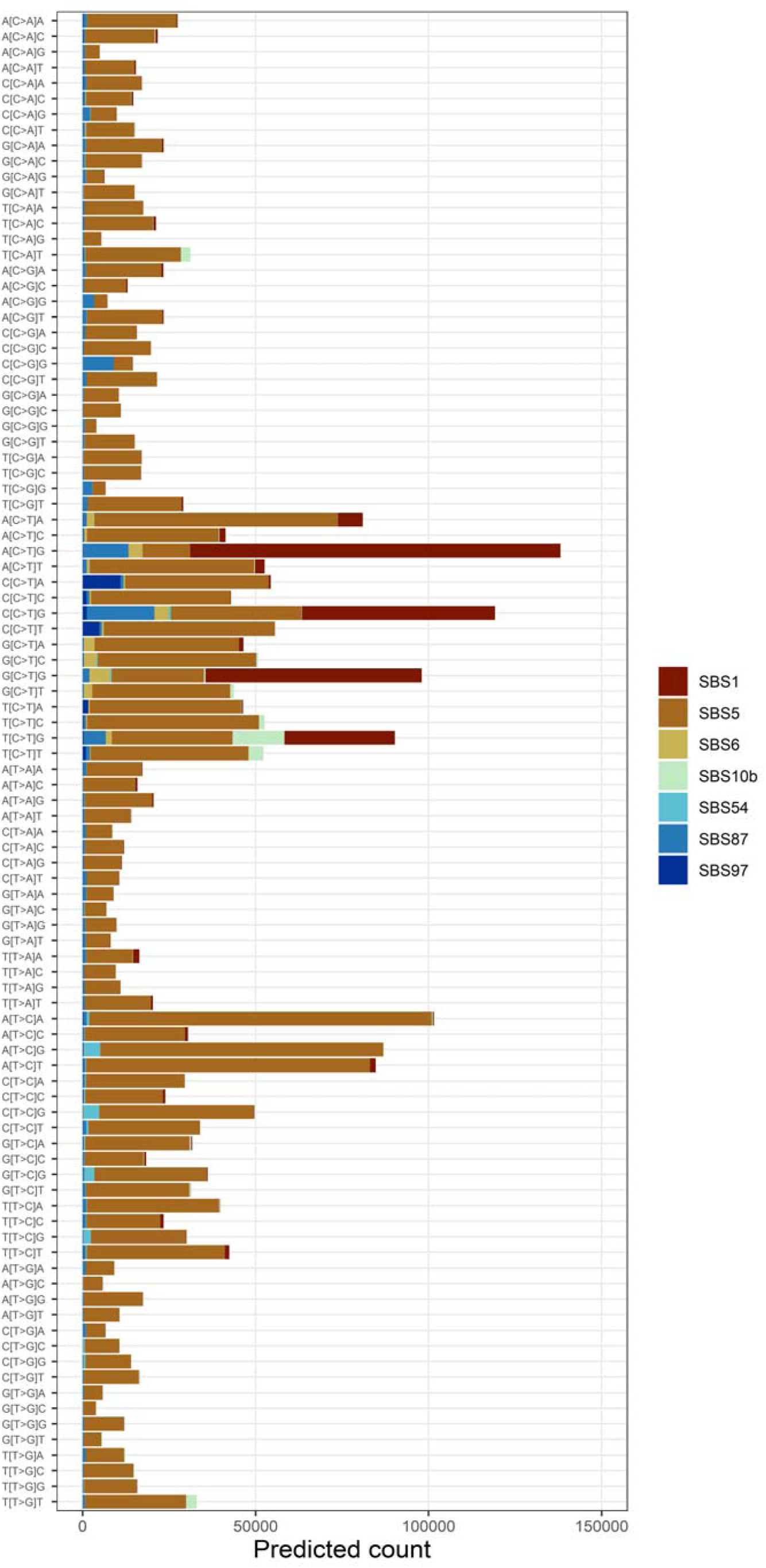
Results of modelling AC25 SBS counts using reference mutational signatures obtained from studies of somatic mutations observed in cancers. **Figure 1a** Observed counts of AC25 SBS variants on trinucleotide backgrounds. Alt text: A horizontal bar chart shows the raw counts of different AC25 variants in the context of their trinucleotide background. **Figure 1b** Predicted AC25 SBS counts using best-fitting combination of seven reference mutational signatures. Alt text: A stacked horizontal bar chart illustrates how seven mutational signatures can approximate the observed counts for AC25 SBS variants occurring on their trinucleotide background.

**Table 1.** Mutational signatures producing the best fit to observed counts of trinucleotide AC25 variants. The mutational signatures are obtained from somatic mutations observed in cancer cells and their proposed aetiologies are as stated for the COSMIC database at https://cancer.sanger.ac.uk/signatures/sbs/.

| Signature | Relative contribution | Proposed aetiology as described in COSMIC database |
| --- | --- | --- |
| SBS1 | 0.178 | Spontaneous or enzymatic deamination of 5-methylcytosine (clock-like signature) |
| SBS5 | 0.486 | Unknown (clock-like signature) |
| SBS6 | 0.061 | Defective DNA mismatch repair |
| SBS10b | 0.061 | Polymerase epsilon exonuclease domain mutations |
| SBS54 | 0.050 | Possible sequencing artefact |
| SBS87 | 0.107 | Thiopurine chemotherapy treatment |
| SBS97 | 0.057 | Unknown |

### Modelling effects of background on variant frequencies

Table 2 shows the frequencies of AC25 variants in the non-transcribed region observed on the background of a C or T nucleotide. It can be seen that the most frequent AC25 variant is C>T, with frequency 0.000919, and the rarest is T>A, with frequency 0.000113. AC25 variant frequencies for backgrounds including different numbers of flanking nucleotides were also calculated. For each partial background sequence, these frequencies could be used to predict the expected observed frequencies for each AC25 variant in the context of the full pentanucleotide background. If just the core nucleotide was used then the predicted frequencies correlated with the observed frequencies with R = 0.508. If a single nucleotide upstream was included the correlation increased to R = 0.746 and including the second upstream nucleotide yielded R = 0.747, while including the two flanking nucleotides yielded R = 0.992. Finally, if two upstream and one downstream nucleotide were included the correlation was R = 0.995. These results suggest that a major driver for the frequency of each variant is the probability of the central base substitution itself but that this probability is strongly influenced by the two neighbouring nucleotides.

**Table 2.** The table shows frequencies for the six different single nucleotide variants having C or T as the background (reference) base relative to the counts of the reference base for AC25 variants in the non-transcribed region of the genome.

| Substitution | Frequencies<br>of AC25<br>variants |
| --- | --- |
| C>A | 0.000256 |
| C>G | 0.000204 |
| C>T | 0.000919 |
| T>A | 0.000113 |
| T>C | 0.000418 |
| T>G | 0.000129 |

Logistic regression modelling was used to better understand which features of the two flanking nucleotides had most effect on the frequency. Figure 2a shows the frequencies of all AC25 variants occurring on trinucleotide sequences with either C or T as the central base. Figure 2b shows the odds ratio (OR) for each trinucleotide sequence to produce the target base relative to the odds for just the central base. The graph does not show the results for variant types having the pattern X[C>T]G because these all had very large ORs compared to the others, with ORs of 8.87, 9.36, 10.17 and 11.16 for T[C>T]G, G[C>T]G, C[C>T]G and A[C>T]G respectively. Among the other variant types, the two for which the flanking nucleotides have the strongest positive effect are A[T>C]A and A[T>C]G with ORs of 2.26 and 2.24 respectively. Conversely T[C>T]T and T[C>T]A have ORs of 0.421 and 0.448, indicating that these flanking nucleotides act to reduce the probability of the variant occurring. Results for all trinucleotide backgrounds having C or T as the central base are presented in Supplementary Table 1.

**Figure 2.**
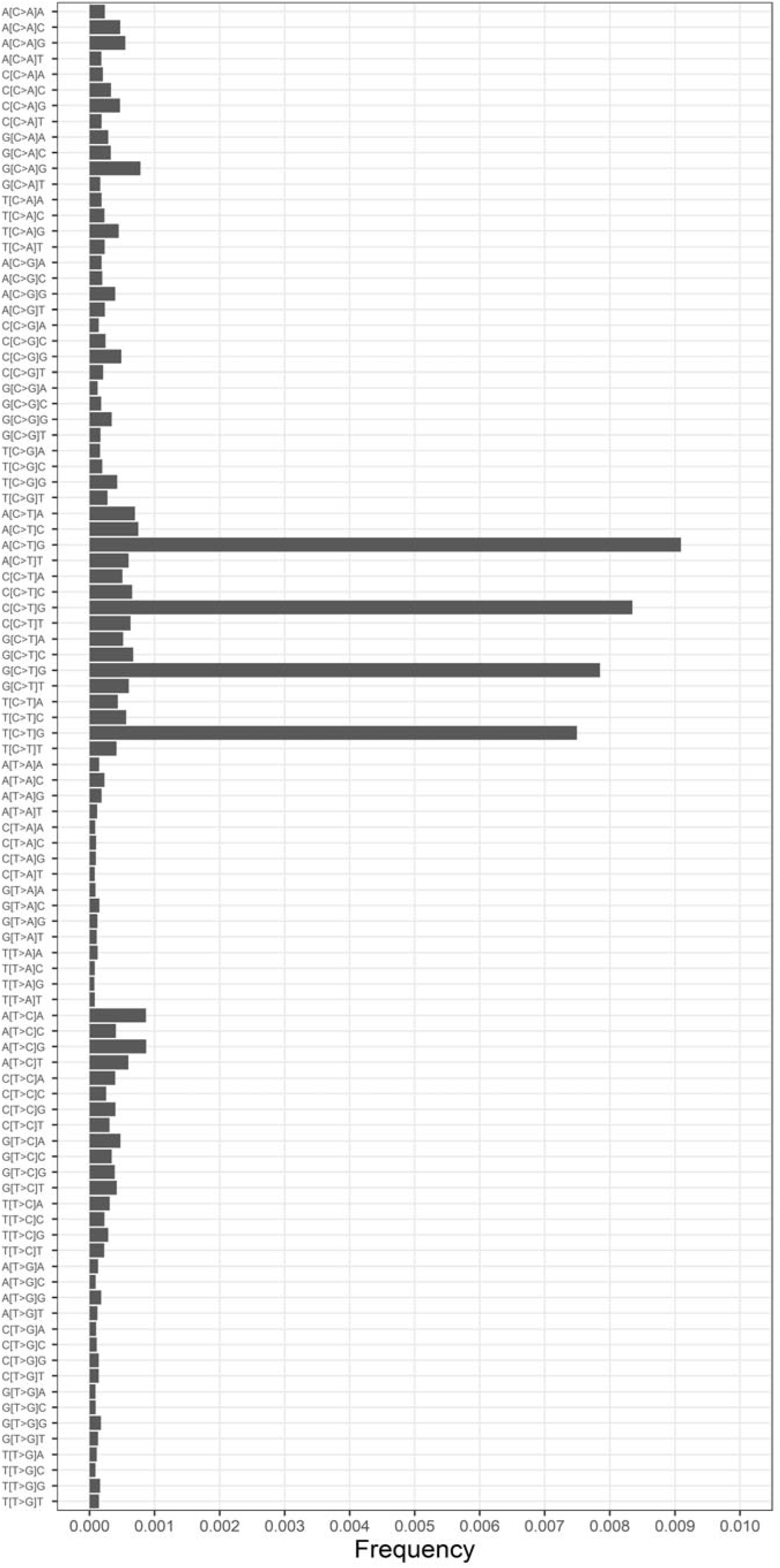
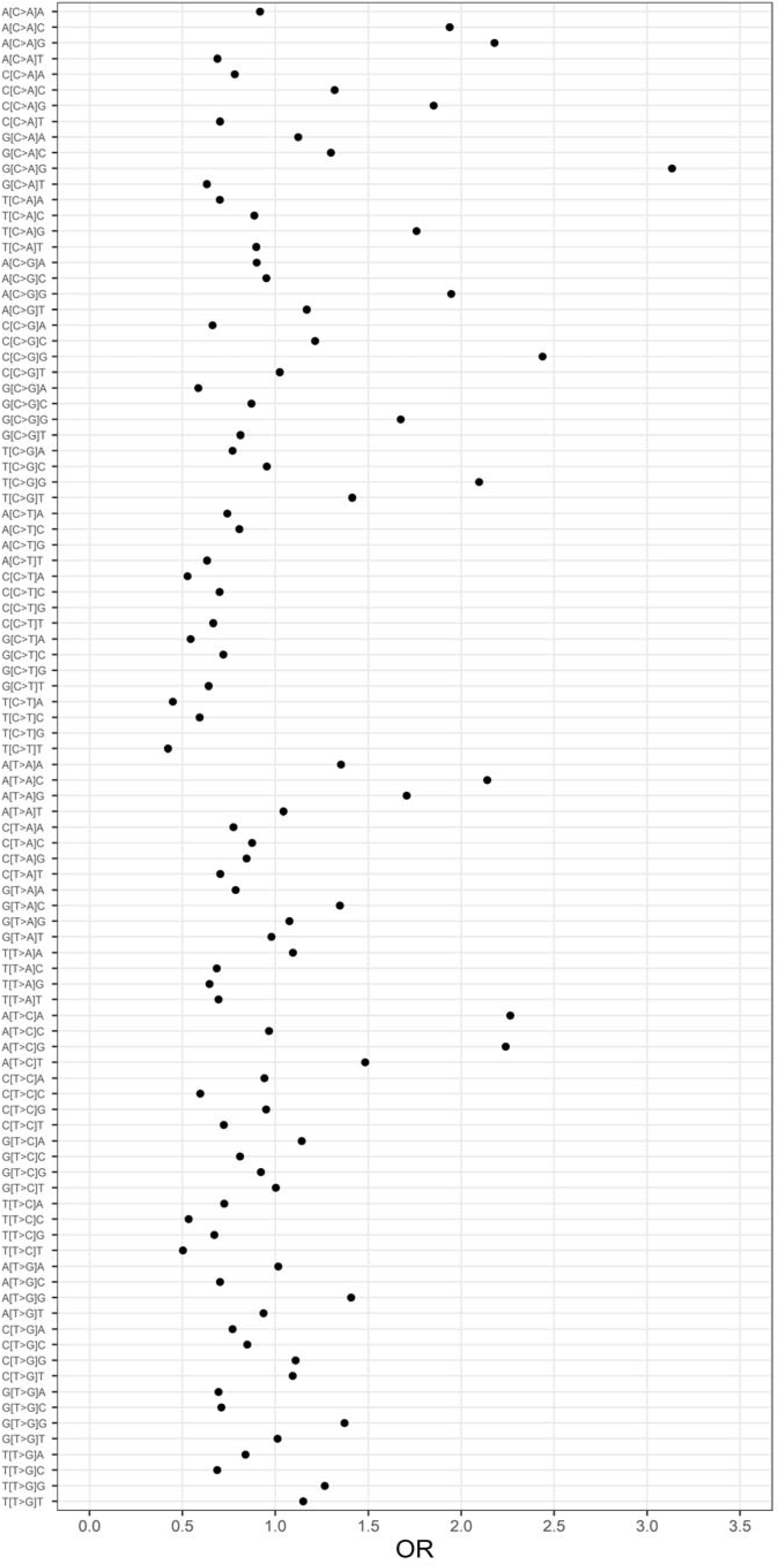
Effects of different trinucleotide sequences on the frequencies of different AC25 variants. Results are shown for all trinucleotides having either C or T as the central base. **Figure 2a** Frequencies of all AC25 variants occurring in the context of different trinucleotide backgrounds. The frequency is defined as the count of the number of times each AC25 variant is observed divided by the count of the number of times the background is seen in the reference genome. Alt text: A horizontal bar chart shows the frequency of each singleton variant with C or T as its central base relative to its trinucleotide background. **Figure 2b** Odds ratio (OR) for each AC25 variant associated with its trinucleotide background, defined as the frequency with which the variant occurs on this background compared with the overall frequency of AC25 single base substitutions of this type (e.g. frequency of A[T>A]C versus frequency of T>A). The results for X[C>T]G variants are not shown because the ORs are too high. Alt text: Each singleton variant on its trinucleotide background is plotted with the horizontal position depicting the odds ratio for the frequency of this variant relative to the central single nucleotide variant irrespective of background.

To evaluate the effects of the full pentanucleotide background compared to the trinucleotide background, the modelling procedure was repeated to obtain the OR for each pentanucleotide sequence to produce the target base relative to the odds of the central trinucleotide sequence. Table 3 shows the variant types with the ten lowest and highest ORs. These results indicate that the more distant nucleotides can have major effects. For example, if there is a C upstream and downstream then the frequency of G[C>A]A and G[C>A]G variants is reduced by a factor of more than 2. Similar results are seen for C[C>G]A variants if there is a T upstream and a C or T downstream. Conversely, TT[T>A]AA and CA[T>C]TG variants have a four-fold higher frequency than the mean for T[T>A]A and A[T>C]T variants. For A[T>C]T variants, having a G downstream and a C, G or T upstream more than doubles the frequency. Results for all pentanucleotide backgrounds having C or T as the central base are presented in Supplementary Table 2.

**Table 3.** Table showing the effect of the full pentanucleotide sequence versus the central trinucleotide sequence on the frequencies of AC25 variants in the non-transcribed region. Results are shown for the ten variants with T or C as the central base for which the frequency is most strongly reduced or increased.

| Variant | OR | (0.95 CI) |
| --- | --- | --- |
| CG[C>A]AC | 0.281 | (0.171 - 0.464) |
| CG[C>A]GC | 0.371 | (0.266 - 0.516) |
| TC[C>G]AC | 0.382 | (0.305 - 0.478) |
| CG[C>A]GA | 0.395 | (0.243 - 0.643) |
| TC[C>A]CA | 0.408 | (0.368 - 0.453) |
| TA[T>A]CG | 0.408 | (0.234 - 0.710) |
| CT[T>A]AG | 0.415 | (0.321 - 0.536) |
| GC[C>A]CG | 0.417 | (0.311 - 0.561) |
| TC[C>G]AT | 0.419 | (0.350 - 0.502) |
| GA[T>A]CG | 0.434 | (0.277 - 0.679) |
| TA[T>C]TG | 2.458 | (2.358 - 2.563) |
| CC[C>G]CC | 2.502 | (2.286 - 2.739) |
| GA[T>G]AG | 2.586 | (2.303 - 2.903) |
| CG[C>G]GA | 2.621 | (1.945 - 3.532) |
| AA[T>C]TG | 2.707 | (2.603 - 2.815) |
| GA[T>C]TG | 2.804 | (2.672 - 2.943) |
| CC[T>G]TC | 2.821 | (2.591 - 3.071) |
| CC[C>A]CC | 3.326 | (3.101 - 3.567) |
| CA[T>C]TG | 3.947 | (3.805 - 4.094) |
| TT[T>A]AA | 4.345 | (4.071 - 4.637) |

### Analysing effects related to transcription

The frequencies of AC25 variants in protein coding transcripts and RNA gene transcripts both correlated with the frequencies for variants in non-transcribed regions with R > 0.995. Nevertheless, the absolute and relative frequencies did differ to some extent. Table 4 displays the differences in frequencies of single nucleotide variants between the different regions. This shows for example that many variants are somewhat less frequent in transcripts of protein coding genes than in the non-transcribed region but that A>G variants are somewhat commoner on the coding strand of protein coding gene transcripts than they are in the non-transcribed region, with OR 1.204 (1.196 - 1.213). However the same variant occurs less commonly on the template strand since the OR for T>C variants is 0.775 (0.768 - 0.781). The same effects can be seen to be present, though less strongly, for transcripts of RNA genes compared to the non-transcribed region, with corresponding ORs of 1.082 (1.071 - 1.093) and 0.907 (0.897 - 0.917). The ORs for these frequencies between RNA gene and protein coding transcripts have narrow confidence intervals, 0.898 (0.889 - 0.908) and 1.171 (1.158 - 1.185), suggesting that these differences are non-random.

**Table 4.**
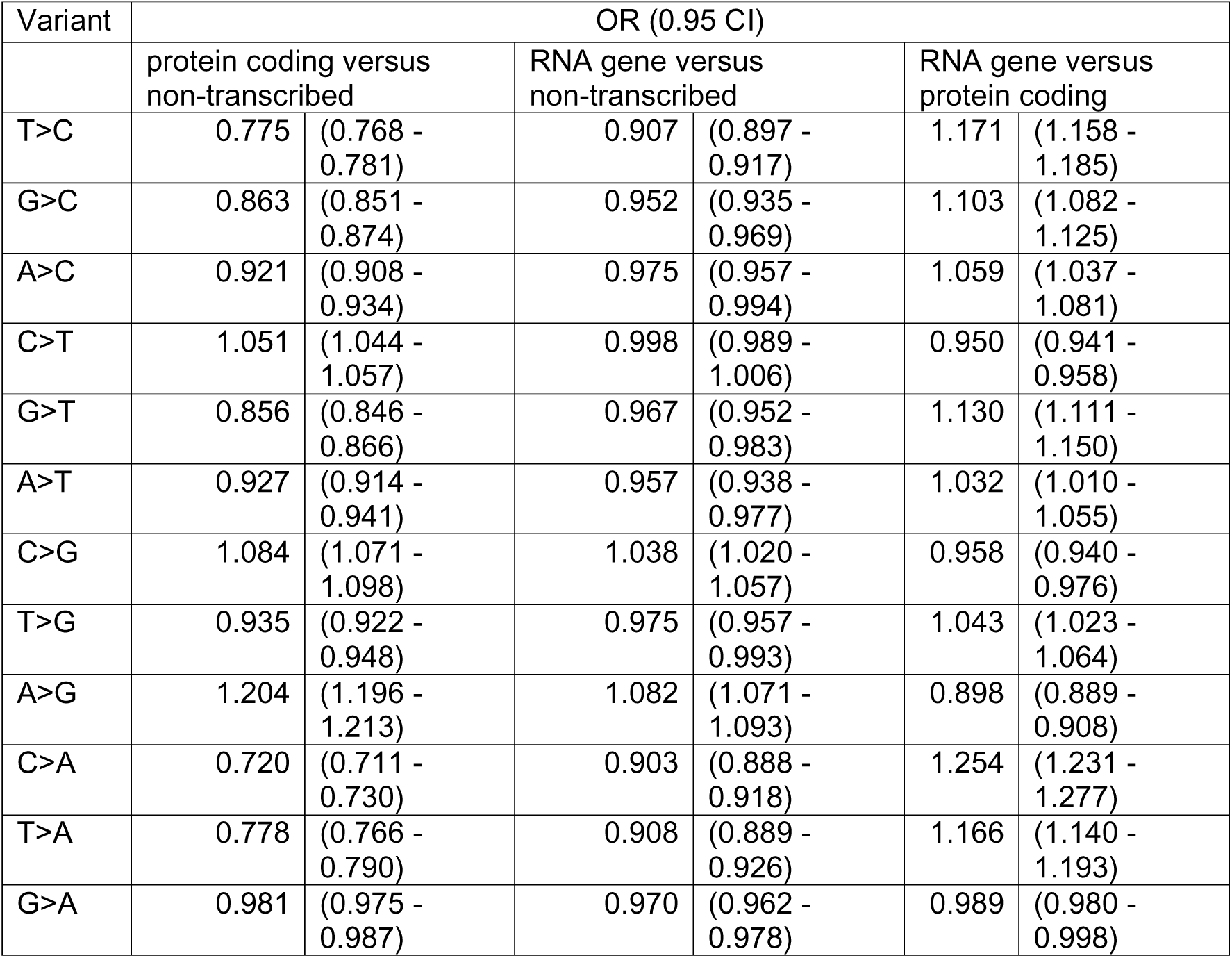
The table shows the ORs of frequencies for AC25 variants between three different regions of the genome, consisting of the non-transcribed region, the transcripts of protein coding genes and the transcripts of RNA genes. For the non-transcribed region variants are defined by the plus strand and for the transcribed region variants are defined by the coding strand.

Similarly to the analyses of effects of sequence context on mutation probability, logistic regression modelling was used to determine how strongly subsets numbers of nucleotides predicted the ORs of frequencies of pentanucleotide variants between different regions. The correlation coefficients for these predictions are presented in Table 5. In general, there are two notable findings. One is that, when predicting ORs relative to non-transcribed regions, the correlation coefficients are lower for the RNA genes than for the protein coding genes. The other is that the correlation coefficients for the protein coding genes are considerably lower than were observed for predicting sequence effects on raw mutation frequencies. For example, the frequencies for the central trinucleotide predicted the raw pentanucleotide frequencies in the non-transcribed region with R = 0.992 but the trinucleotide ORs for frequency differences between the non-transcribed and protein coding region predict the observed pentanucleotide ORs with R = 0.602. An interpretation of this might be that the relative probability for a mutation to occur in the non-transcribed region versus a protein coding transcript is not strongly determined by the immediately flanking nucleotides, as defined by the trinucleotide, but is more dependent on the full pentanucleotide context.

**Table 5.** The table shows the correlation coefficient between the ORs of observed frequencies of full pentanucleotide AC25 variants between regions and the ORs predicted from using a subset of the component nucleotides.

| Nucleotides included | Correlation coefficient between predicted and actual pentanucleotide variant frequency ORs |  |  |
| --- | --- | --- | --- |
|  | protein coding versus non-transcribed | RNA gene versus non-transcribed | RNA gene versus protein coding |
| Central nucleotide only | 0.536 | 0.200 | 0.312 |
| Central and one upstream | 0.581 | 0.228 | 0.355 |
| Central and one downstream | 0.580 | 0.221 | 0.348 |
| Central and two upstream | 0.643 | 0.287 | 0.414 |
| Central and two downstream | 0.630 | 0.294 | 0.416 |
| Central and two flanking | 0.602 | 0.279 | 0.393 |
| Central, two upstream, one downstream | 0.744 | 0.491 | 0.564 |
| Central, one upstream, two downstream | 0.713 | 0.485 | 0.566 |

Table 6 shows the twenty variants which have the most significantly increased frequencies in protein coding and RNA gene transcripts compared to the non-transcribed region, as assessed by having the largest values for z = ln(OR)/SE which under the null hypothesis follows a gaussian distribution with zero mean and unit standard deviation. Strikingly, it can be seen that the eight variants with the strongest evidence for increased frequencies in transcripts of protein coding genes all have the motif CT[A>G]T* or CA[A>G]T* and these all have OR > 1.5. All of the top twenty variants have the A>G substitution. For the RNA gene transcripts effect sizes are smaller and less strongly statistically significant but it seems that similar effects operate in that the top four variants match one of these motifs and all but one of the top twenty has the A>G substitution. To explore whether there were systematic differences between the protein coding gene transcripts and the RNA gene transcripts, their variant frequencies were compared directly and the results are shown in Table 7. This shows that, for example, variants matching the CT[A>G]T* and CA[A>G]T* are significantly more frequent in protein coding gene transcripts than in RNA gene transcripts. Conversely there are variants which are more common in RNA gene transcripts than protein coding gene transcripts and these include some with motifs *A[T>C]AG and *A[T>C]TG. Notably, these motifs are complementary to the CT[A>G]T* and CA[A>G]T* motifs, meaning that another way to express these findings would be to say that CT[A>G]T* and CA[A>G]T* variants on the coding strand are more frequent in protein coding than RNA gene transcripts but on the template strand they are more frequent in RNA gene than protein coding gene transcripts.

**Table 6.**
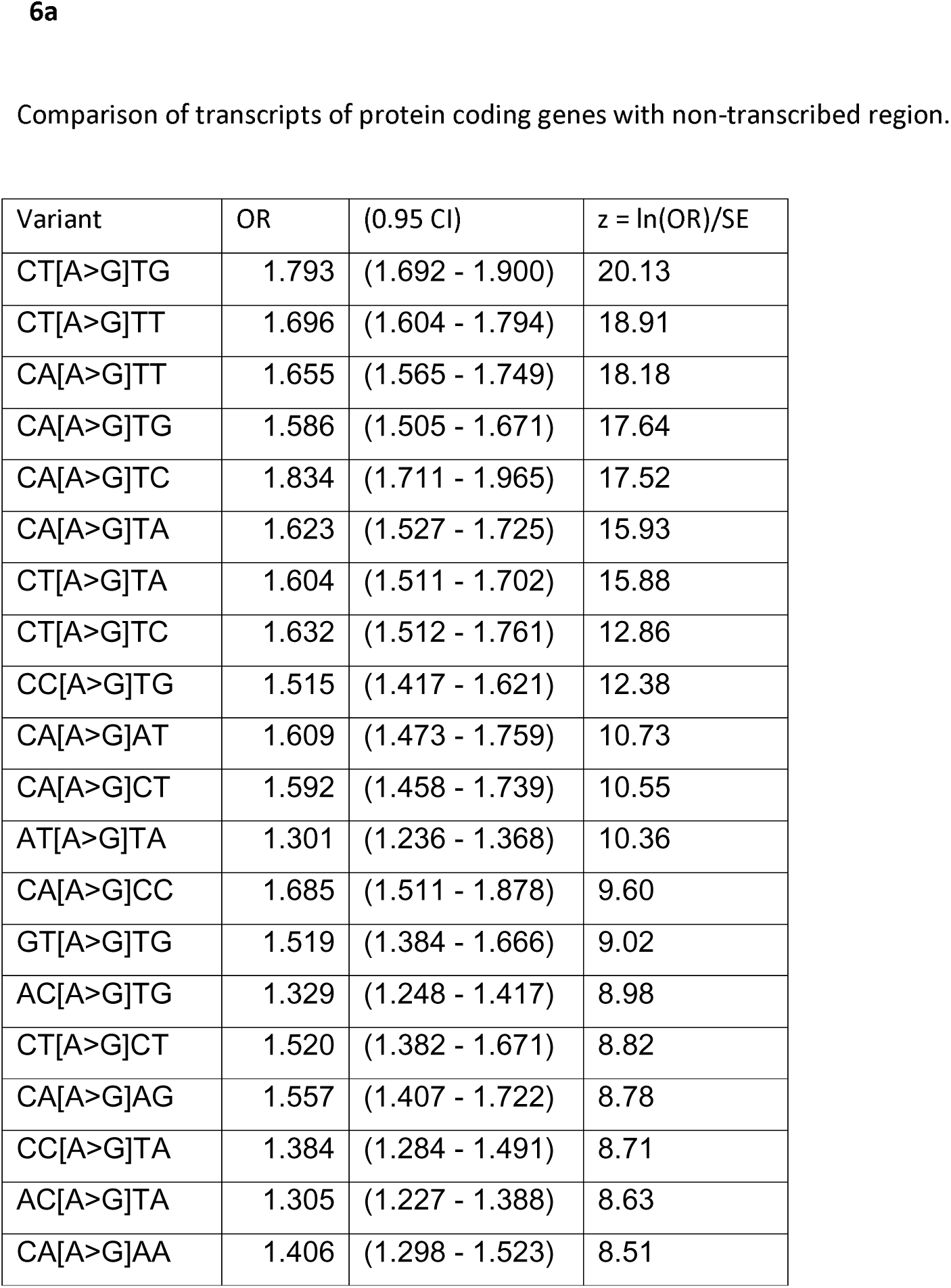
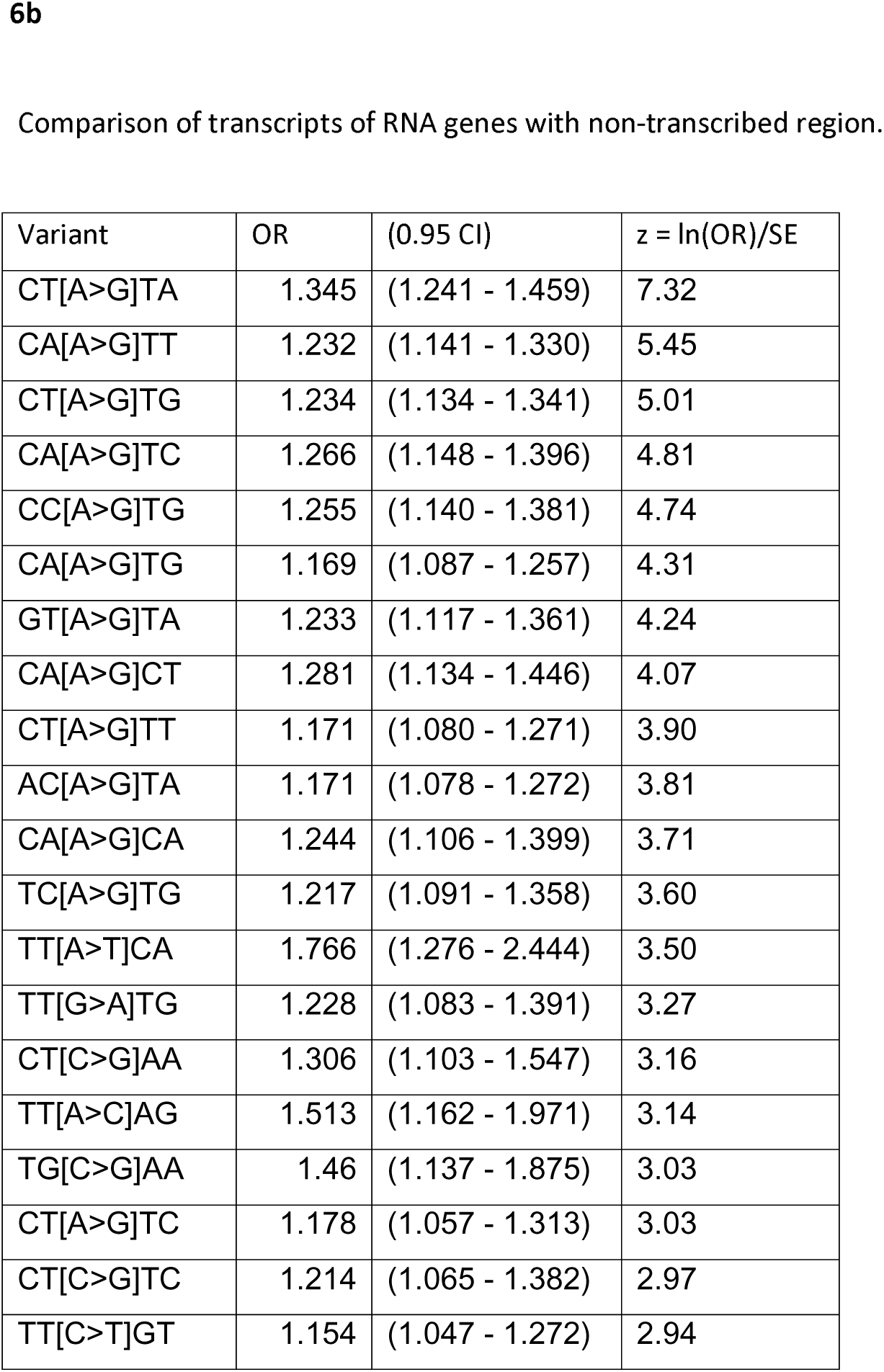
Table showing the variants with the strongest evidence for having a higher frequency in transcripts of protein coding genes and RNA genes relative to their frequency in the non-transcribed region.

| Variant | OR | (0.95 CI) | $z = \ln(OR)/SE$ |
| --- | --- | --- | --- |
| CT[A>G]TG | 1.793 | (1.692 - 1.900) | 20.13 |
| CT[A>G]TT | 1.696 | (1.604 - 1.794) | 18.91 |
| CA[A>G]TT | 1.655 | (1.565 - 1.749) | 18.18 |
| CA[A>G]TG | 1.586 | (1.505 - 1.671) | 17.64 |
| CA[A>G]TC | 1.834 | (1.711 - 1.965) | 17.52 |
| CA[A>G]TA | 1.623 | (1.527 - 1.725) | 15.93 |
| CT[A>G]TA | 1.604 | (1.511 - 1.702) | 15.88 |
| CT[A>G]TC | 1.632 | (1.512 - 1.761) | 12.86 |
| CC[A>G]TG | 1.515 | (1.417 - 1.621) | 12.38 |
| CA[A>G]AT | 1.609 | (1.473 - 1.759) | 10.73 |
| CA[A>G]CT | 1.592 | (1.458 - 1.739) | 10.55 |
| AT[A>G]TA | 1.301 | (1.236 - 1.368) | 10.36 |
| CA[A>G]CC | 1.685 | (1.511 - 1.878) | 9.60 |
| GT[A>G]TG | 1.519 | (1.384 - 1.666) | 9.02 |
| AC[A>G]TG | 1.329 | (1.248 - 1.417) | 8.98 |
| CT[A>G]CT | 1.520 | (1.382 - 1.671) | 8.82 |
| CA[A>G]AG | 1.557 | (1.407 - 1.722) | 8.78 |
| CC[A>G]TA | 1.384 | (1.284 - 1.491) | 8.71 |
| AC[A>G]TA | 1.305 | (1.227 - 1.388) | 8.63 |
| CA[A>G]AA | 1.406 | (1.298 - 1.523) | 8.51 |
| CT[A>G]TA | 1.345 | (1.241 - 1.459) | 7.32 |
| CA[A>G]TT | 1.232 | (1.141 - 1.330) | 5.45 |
| CT[A>G]TG | 1.234 | (1.134 - 1.341) | 5.01 |
| CA[A>G]TC | 1.266 | (1.148 - 1.396) | 4.81 |
| CC[A>G]TG | 1.255 | (1.140 - 1.381) | 4.74 |
| CA[A>G]TG | 1.169 | (1.087 - 1.257) | 4.31 |
| GT[A>G]TA | 1.233 | (1.117 - 1.361) | 4.24 |
| CA[A>G]CT | 1.281 | (1.134 - 1.446) | 4.07 |
| CT[A>G]TT | 1.171 | (1.080 - 1.271) | 3.90 |
| AC[A>G]TA | 1.171 | (1.078 - 1.272) | 3.81 |
| CA[A>G]CA | 1.244 | (1.106 - 1.399) | 3.71 |
| TC[A>G]TG | 1.217 | (1.091 - 1.358) | 3.60 |
| TT[A>T]CA | 1.766 | (1.276 - 2.444) | 3.50 |
| TT[G>A]TG | 1.228 | (1.083 - 1.391) | 3.27 |
| CT[C>G]AA | 1.306 | (1.103 - 1.547) | 3.16 |
| TT[A>C]AG | 1.513 | (1.162 - 1.971) | 3.14 |
| TG[C>G]AA | 1.46 | (1.137 - 1.875) | 3.03 |
| CT[A>G]TC | 1.178 | (1.057 - 1.313) | 3.03 |
| CT[C>G]TC | 1.214 | (1.065 - 1.382) | 2.97 |
| TT[C>T]GT | 1.154 | (1.047 - 1.272) | 2.94 |

**Table 7.**
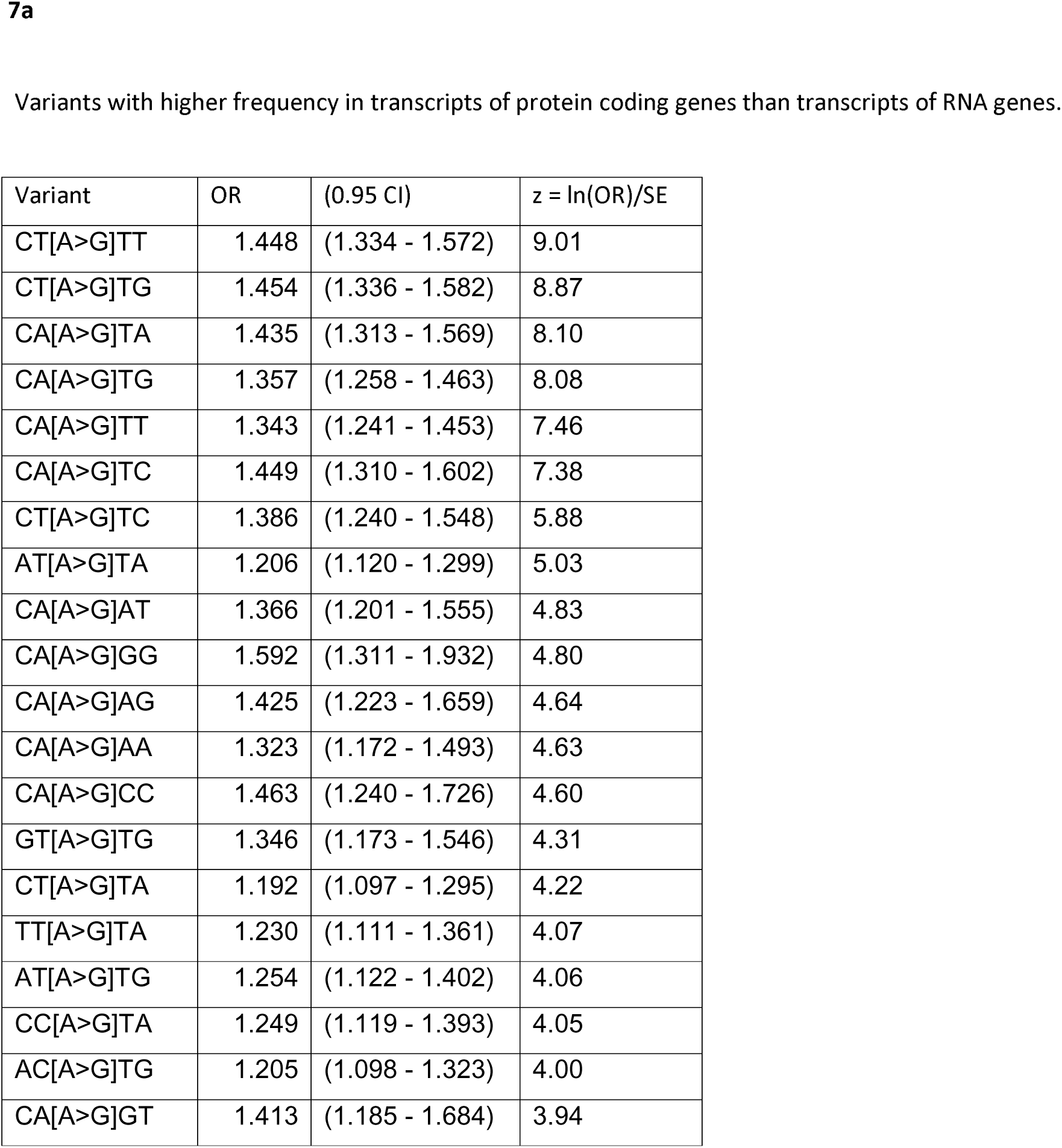
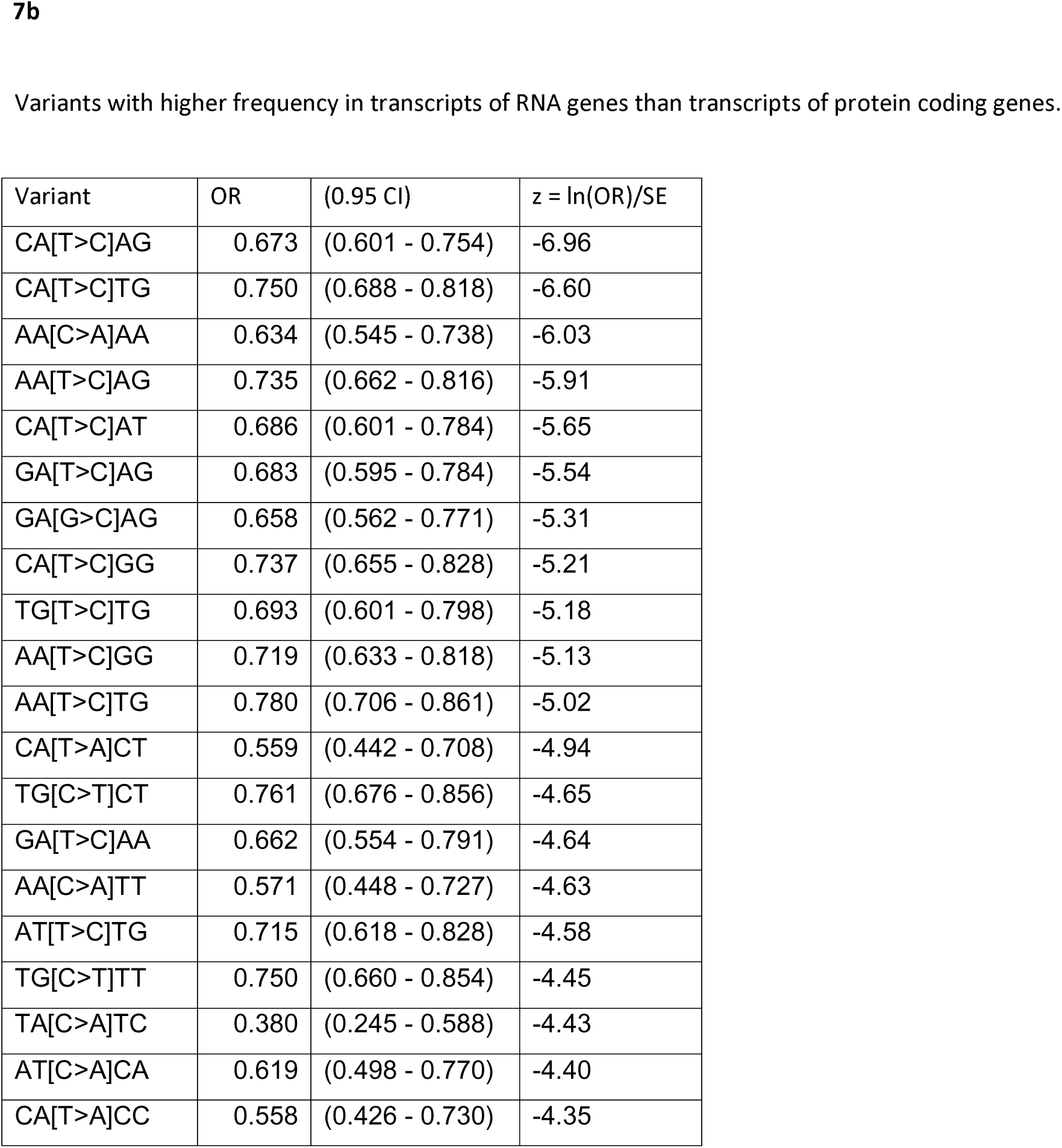
Table showing the variants with the strongest evidence for having a higher frequency in transcripts of protein coding genes relative to their frequency in transcripts of RNA genes or vice versa.

| Variant | OR | (0.95 CI) | $z = \ln(OR)/SE$ |
| --- | --- | --- | --- |
| CT[A>G]TT | 1.448 | (1.334 - 1.572) | 9.01 |
| CT[A>G]TG | 1.454 | (1.336 - 1.582) | 8.87 |
| CA[A>G]TA | 1.435 | (1.313 - 1.569) | 8.10 |
| CA[A>G]TG | 1.357 | (1.258 - 1.463) | 8.08 |
| CA[A>G]TT | 1.343 | (1.241 - 1.453) | 7.46 |
| CA[A>G]TC | 1.449 | (1.310 - 1.602) | 7.38 |
| CT[A>G]TC | 1.386 | (1.240 - 1.548) | 5.88 |
| AT[A>G]TA | 1.206 | (1.120 - 1.299) | 5.03 |
| CA[A>G]AT | 1.366 | (1.201 - 1.555) | 4.83 |
| CA[A>G]GG | 1.592 | (1.311 - 1.932) | 4.80 |
| CA[A>G]AG | 1.425 | (1.223 - 1.659) | 4.64 |
| CA[A>G]AA | 1.323 | (1.172 - 1.493) | 4.63 |
| CA[A>G]CC | 1.463 | (1.240 - 1.726) | 4.60 |
| GT[A>G]TG | 1.346 | (1.173 - 1.546) | 4.31 |
| CT[A>G]TA | 1.192 | (1.097 - 1.295) | 4.22 |
| TT[A>G]TA | 1.230 | (1.111 - 1.361) | 4.07 |
| AT[A>G]TG | 1.254 | (1.122 - 1.402) | 4.06 |
| CC[A>G]TA | 1.249 | (1.119 - 1.393) | 4.05 |
| AC[A>G]TG | 1.205 | (1.098 - 1.323) | 4.00 |
| CA[A>G]GT | 1.413 | (1.185 - 1.684) | 3.94 |
| CA[T>C]AG | 0.673 | (0.601 - 0.754) | -6.96 |
| CA[T>C]TG | 0.750 | (0.688 - 0.818) | -6.60 |
| AA[C>A]AA | 0.634 | (0.545 - 0.738) | -6.03 |
| AA[T>C]AG | 0.735 | (0.662 - 0.816) | -5.91 |
| CA[T>C]AT | 0.686 | (0.601 - 0.784) | -5.65 |
| GA[T>C]AG | 0.683 | (0.595 - 0.784) | -5.54 |
| GA[G>C]AG | 0.658 | (0.562 - 0.771) | -5.31 |
| CA[T>C]GG | 0.737 | (0.655 - 0.828) | -5.21 |
| TG[T>C]TG | 0.693 | (0.601 - 0.798) | -5.18 |
| AA[T>C]GG | 0.719 | (0.633 - 0.818) | -5.13 |
| AA[T>C]TG | 0.780 | (0.706 - 0.861) | -5.02 |
| CA[T>A]CT | 0.559 | (0.442 - 0.708) | -4.94 |
| TG[C>T]CT | 0.761 | (0.676 - 0.856) | -4.65 |
| GA[T>C]AA | 0.662 | (0.554 - 0.791) | -4.64 |
| AA[C>A]TT | 0.571 | (0.448 - 0.727) | -4.63 |
| AT[T>C]TG | 0.715 | (0.618 - 0.828) | -4.58 |
| TG[C>T]TT | 0.750 | (0.660 - 0.854) | -4.45 |
| TA[C>A]TC | 0.380 | (0.245 - 0.588) | -4.43 |
| AT[C>A]CA | 0.619 | (0.498 - 0.770) | -4.40 |
| CA[T>A]CC | 0.558 | (0.426 - 0.730) | -4.35 |

Strand asymmetry of variants was assessed directly in the three regions by comparing the frequency of each variant against the complementary sequence or, putting it another way, the frequency of the variant observed relative to the other strand. Results for the variants showing the strongest evidence for asymmetry are shown in Table 8. In the non-transcribed region effects are small, non-systematic and mostly not statistically significant. By contrast, in the transcripts of protein coding genes the CT[A>G]T* and CA[A>G]T* variants all occur more commonly on the coding strand than template strand with ORs exceeding 2. Other variants also exhibit marked asymmetry and all those in the top 20 have the A>G substitution. For the transcripts of RNA genes the effects seem similar though less marked.

**Table 8.**
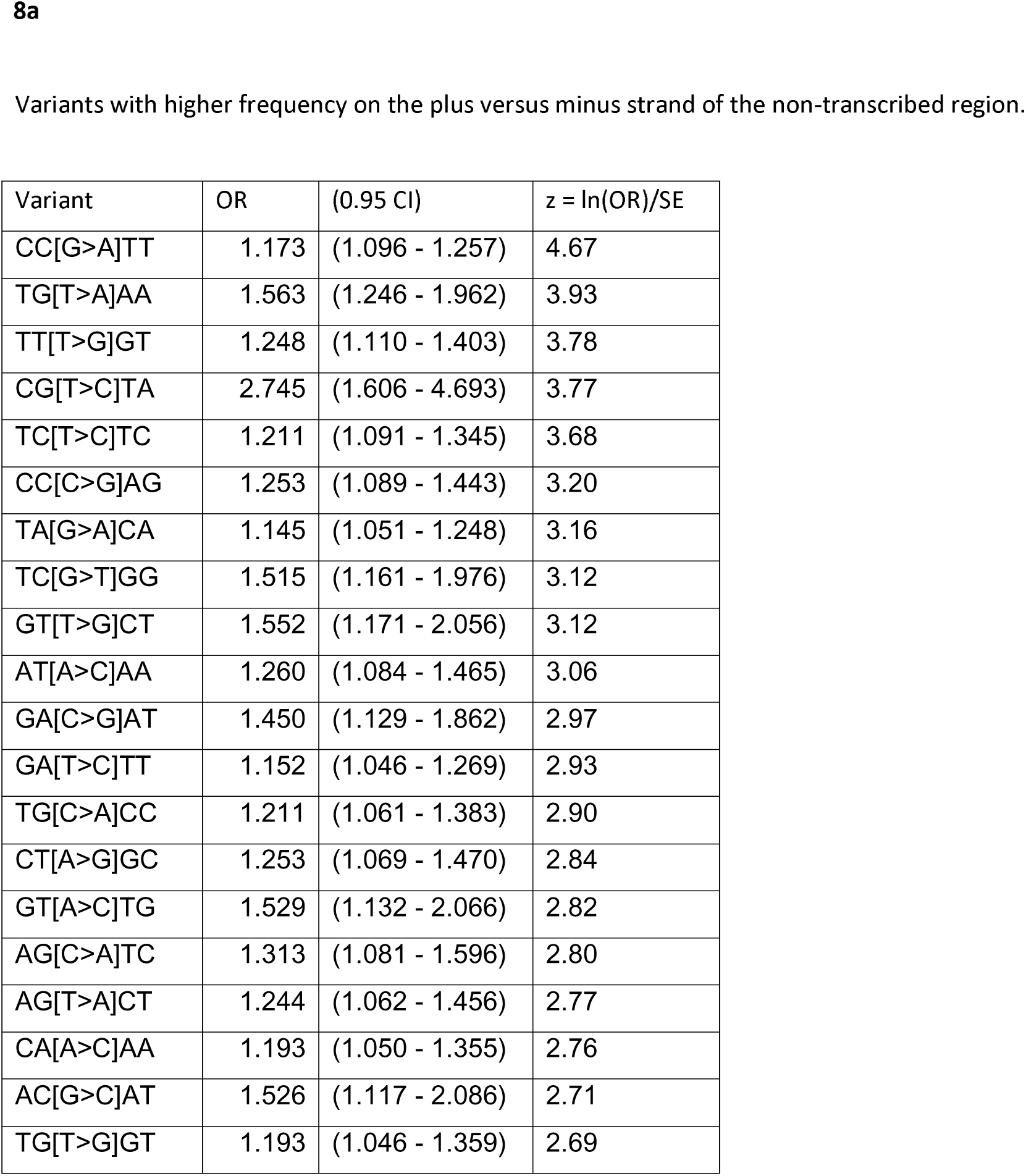
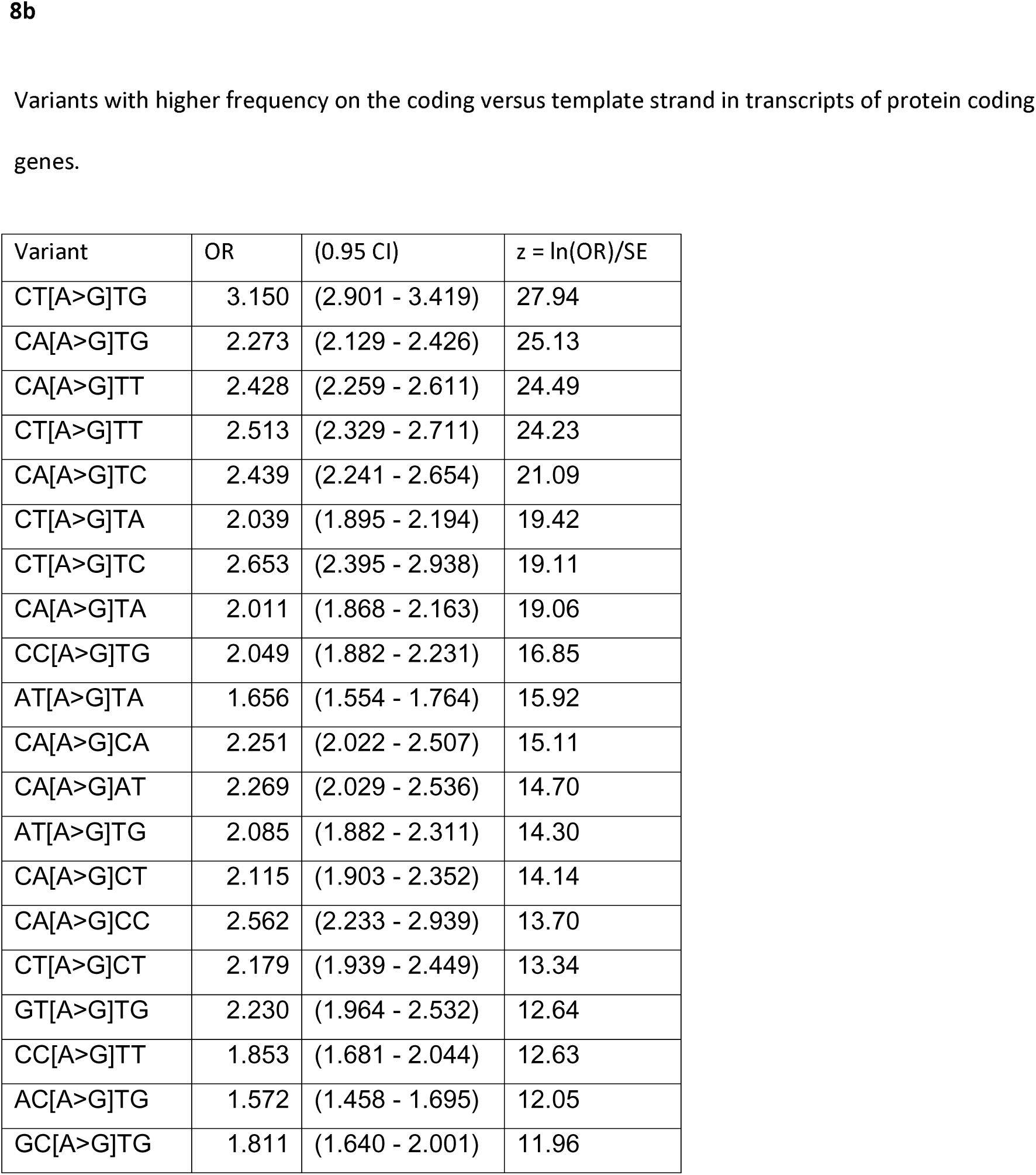
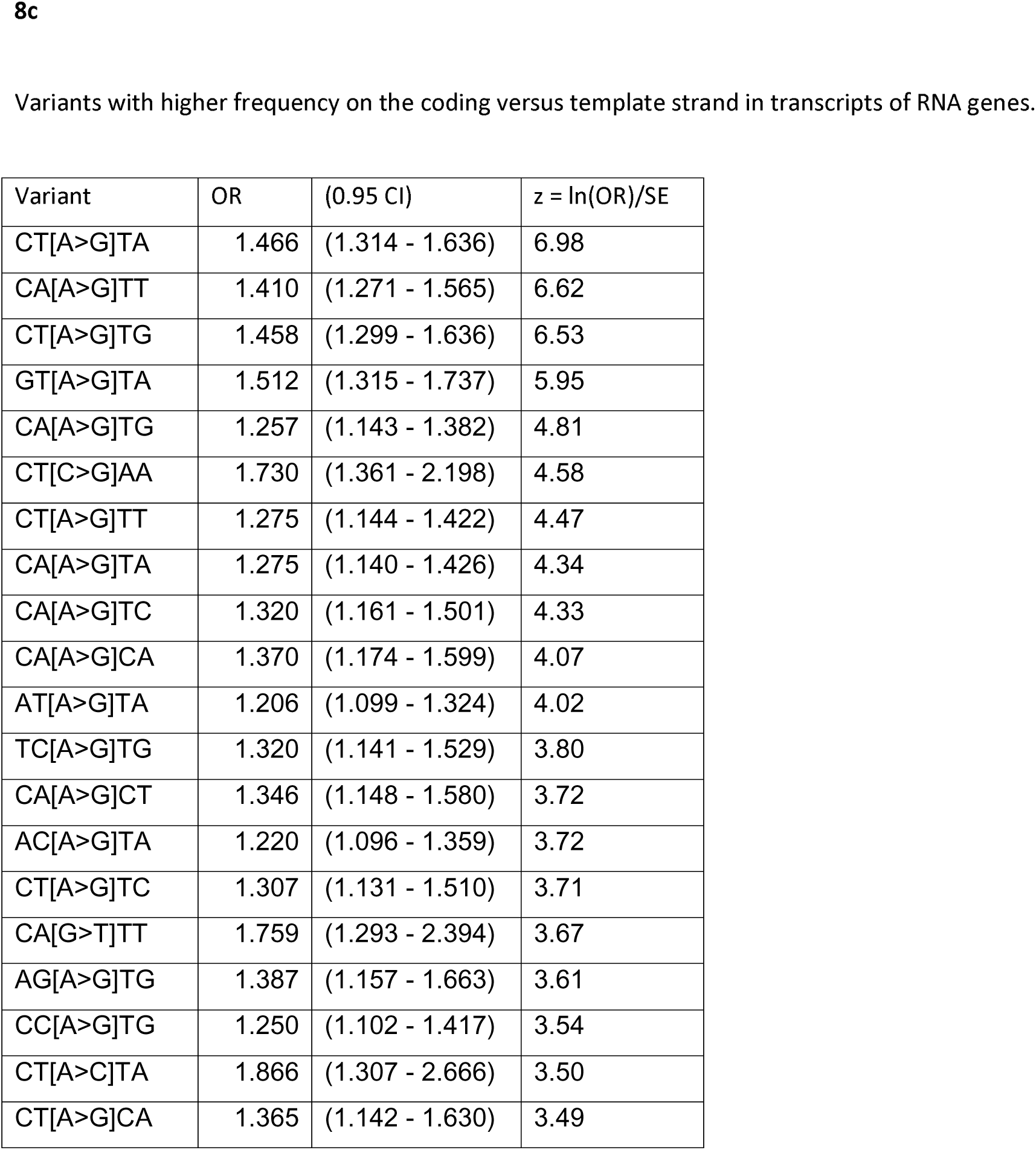
Tables showing the variants with the strongest evidence for strand asymmetry.

| Variant | OR | (0.95 CI) | $z = \ln(OR)/SE$ |
| --- | --- | --- | --- |
| CC[G>A]TT | 1.173 | (1.096 - 1.257) | 4.67 |
| TG[T>A]AA | 1.563 | (1.246 - 1.962) | 3.93 |
| TT[T>G]GT | 1.248 | (1.110 - 1.403) | 3.78 |
| CG[T>C]TA | 2.745 | (1.606 - 4.693) | 3.77 |
| TC[T>C]TC | 1.211 | (1.091 - 1.345) | 3.68 |
| CC[C>G]AG | 1.253 | (1.089 - 1.443) | 3.20 |
| TA[G>A]CA | 1.145 | (1.051 - 1.248) | 3.16 |
| TC[G>T]GG | 1.515 | (1.161 - 1.976) | 3.12 |
| GT[T>G]CT | 1.552 | (1.171 - 2.056) | 3.12 |
| AT[A>C]AA | 1.260 | (1.084 - 1.465) | 3.06 |
| GA[C>G]AT | 1.450 | (1.129 - 1.862) | 2.97 |
| GA[T>C]TT | 1.152 | (1.046 - 1.269) | 2.93 |
| TG[C>A]CC | 1.211 | (1.061 - 1.383) | 2.90 |
| CT[A>G]GC | 1.253 | (1.069 - 1.470) | 2.84 |
| GT[A>C]TG | 1.529 | (1.132 - 2.066) | 2.82 |
| AG[C>A]TC | 1.313 | (1.081 - 1.596) | 2.80 |
| AG[T>A]CT | 1.244 | (1.062 - 1.456) | 2.77 |
| CA[A>C]AA | 1.193 | (1.050 - 1.355) | 2.76 |
| AC[G>C]AT | 1.526 | (1.117 - 2.086) | 2.71 |
| TG[T>G]GT | 1.193 | (1.046 - 1.359) | 2.69 |
| CT[A>G]TG | 3.150 | (2.901 - 3.419) | 27.94 |
| CA[A>G]TG | 2.273 | (2.129 - 2.426) | 25.13 |
| CA[A>G]TT | 2.428 | (2.259 - 2.611) | 24.49 |
| CT[A>G]TT | 2.513 | (2.329 - 2.711) | 24.23 |
| CA[A>G]TC | 2.439 | (2.241 - 2.654) | 21.09 |
| CT[A>G]TA | 2.039 | (1.895 - 2.194) | 19.42 |
| CT[A>G]TC | 2.653 | (2.395 - 2.938) | 19.11 |
| CA[A>G]TA | 2.011 | (1.868 - 2.163) | 19.06 |
| CC[A>G]TG | 2.049 | (1.882 - 2.231) | 16.85 |
| AT[A>G]TA | 1.656 | (1.554 - 1.764) | 15.92 |
| CA[A>G]CA | 2.251 | (2.022 - 2.507) | 15.11 |
| CA[A>G]AT | 2.269 | (2.029 - 2.536) | 14.70 |
| AT[A>G]TG | 2.085 | (1.882 - 2.311) | 14.30 |
| CA[A>G]CT | 2.115 | (1.903 - 2.352) | 14.14 |
| CA[A>G]CC | 2.562 | (2.233 - 2.939) | 13.70 |
| CT[A>G]CT | 2.179 | (1.939 - 2.449) | 13.34 |
| GT[A>G]TG | 2.230 | (1.964 - 2.532) | 12.64 |
| CC[A>G]TT | 1.853 | (1.681 - 2.044) | 12.63 |
| AC[A>G]TG | 1.572 | (1.458 - 1.695) | 12.05 |
| GC[A>G]TG | 1.811 | (1.640 - 2.001) | 11.96 |
| CT[A>G]TA | 1.466 | (1.314 - 1.636) | 6.98 |
| CA[A>G]TT | 1.410 | (1.271 - 1.565) | 6.62 |
| CT[A>G]TG | 1.458 | (1.299 - 1.636) | 6.53 |
| GT[A>G]TA | 1.512 | (1.315 - 1.737) | 5.95 |
| CA[A>G]TG | 1.257 | (1.143 - 1.382) | 4.81 |
| CT[C>G]AA | 1.730 | (1.361 - 2.198) | 4.58 |
| CT[A>G]TT | 1.275 | (1.144 - 1.422) | 4.47 |
| CA[A>G]TA | 1.275 | (1.140 - 1.426) | 4.34 |
| CA[A>G]TC | 1.320 | (1.161 - 1.501) | 4.33 |
| CA[A>G]CA | 1.370 | (1.174 - 1.599) | 4.07 |
| AT[A>G]TA | 1.206 | (1.099 - 1.324) | 4.02 |
| TC[A>G]TG | 1.320 | (1.141 - 1.529) | 3.80 |
| CA[A>G]CT | 1.346 | (1.148 - 1.580) | 3.72 |
| AC[A>G]TA | 1.220 | (1.096 - 1.359) | 3.72 |
| CT[A>G]TC | 1.307 | (1.131 - 1.510) | 3.71 |
| CA[G>T]TT | 1.759 | (1.293 - 2.394) | 3.67 |
| AG[A>G]TG | 1.387 | (1.157 - 1.663) | 3.61 |
| CC[A>G]TG | 1.250 | (1.102 - 1.417) | 3.54 |
| CT[A>C]TA | 1.866 | (1.307 - 2.666) | 3.50 |
| CT[A>G]CA | 1.365 | (1.142 - 1.630) | 3.49 |

The detailed results for all comparisons of variant frequencies between regions and between strands are provided in Supplementary Tables 3 and 4.

In order to investigate which genes contributed to these effects, the frequencies of all CT[A>G]T* variants (not just those with allele count = 25) were calculated for each protein coding gene with length >= 15 kb. The 500 genes with the highest and lowest frequencies for these variants were entered into a PAN-GO enrichment analysis for biological pathways and the results are displayed in Table 9. Table 9a shows that the genes with the highest frequencies of these variants tend to be involved in gene expression and RNA metabolism whereas Table 9b shows that genes with the lowest variant frequencies are less likely to be involved in these processes. Instead, Table 9b shows that the genes with the lowest variant frequencies demonstrate enrichment for more specialised processes such as synaptic transmission and muscle contraction.

**Table 9.**
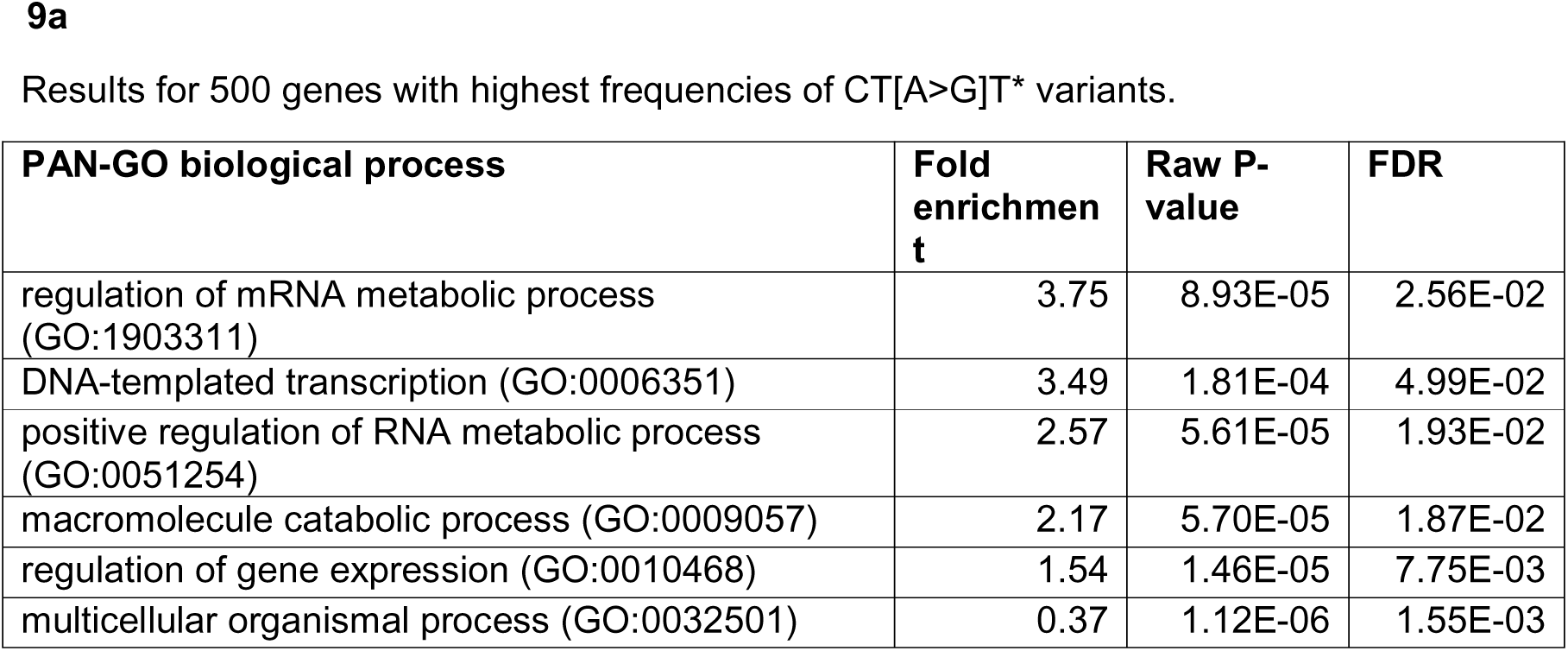
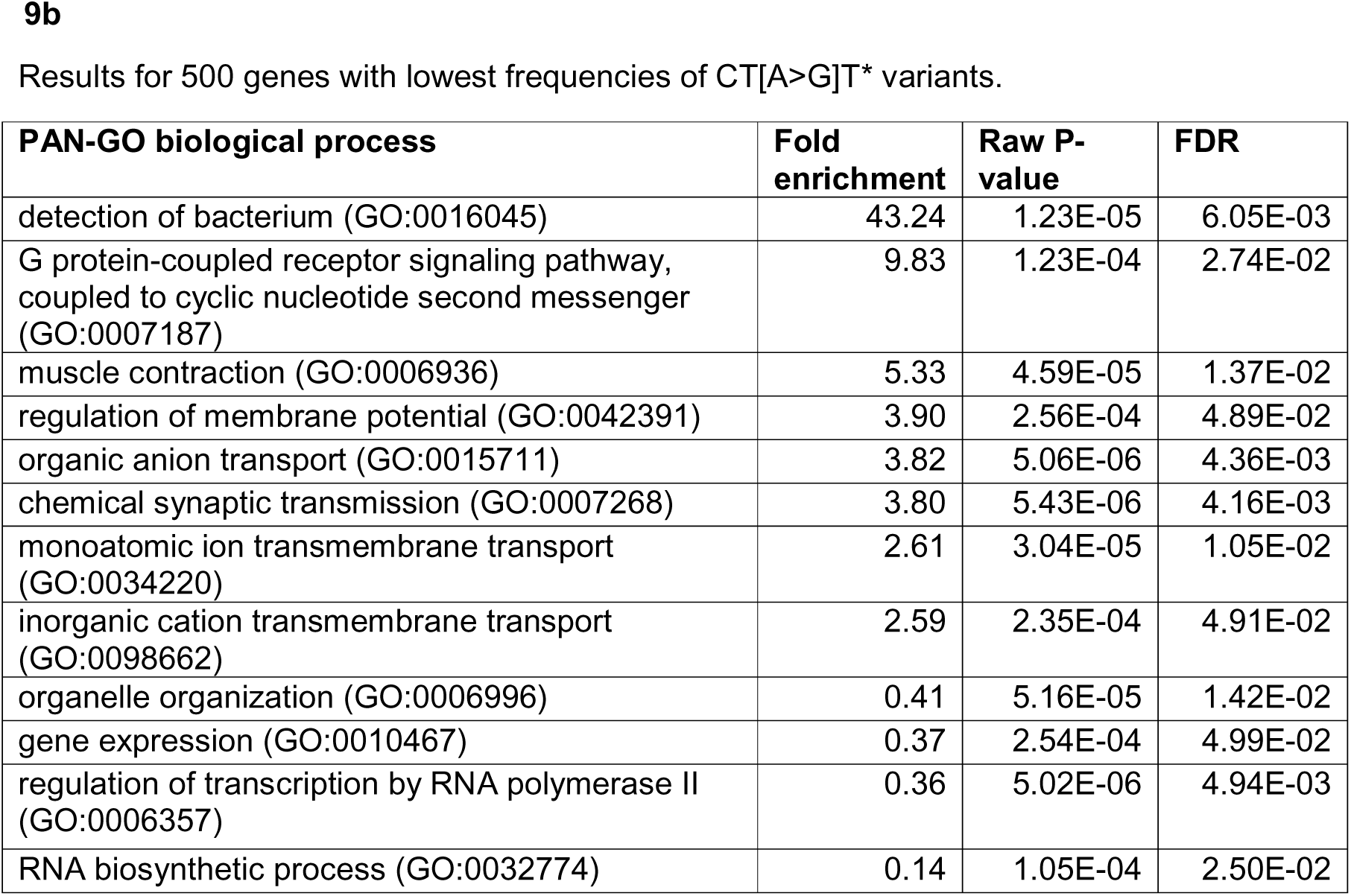
Results of PAN-GO enrichment analyses for genes with the highest and lowest frequencies of CT[A>G]T* variants showing the top level biological processes with statistically significant enrichment at false discovery rate (FDR) < 0.05. A “Fold enrichment” value less than 1 indicates a depletion of genes involved in the relevant pathway.

| PAN-GO biological process | Fold enrichment | Raw P-value | FDR |
| --- | --- | --- | --- |
| regulation of mRNA metabolic process (GO:1903311) | 3.75 | 8.93E-05 | 2.56E-02 |
| DNA-templated transcription (GO:0006351) | 3.49 | 1.81E-04 | 4.99E-02 |
| positive regulation of RNA metabolic process (GO:0051254) | 2.57 | 5.61E-05 | 1.93E-02 |
| macromolecule catabolic process (GO:0009057) | 2.17 | 5.70E-05 | 1.87E-02 |
| regulation of gene expression (GO:0010468) | 1.54 | 1.46E-05 | 7.75E-03 |
| multicellular organismal process (GO:0032501) | 0.37 | 1.12E-06 | 1.55E-03 |
| detection of bacterium (GO:0016045) | 43.24 | 1.23E-05 | 6.05E-03 |
| G protein-coupled receptor signaling pathway, coupled to cyclic nucleotide second messenger (GO:0007187) | 9.83 | 1.23E-04 | 2.74E-02 |
| muscle contraction (GO:0006936) | 5.33 | 4.59E-05 | 1.37E-02 |
| regulation of membrane potential (GO:0042391) | 3.90 | 2.56E-04 | 4.89E-02 |
| organic anion transport (GO:0015711) | 3.82 | 5.06E-06 | 4.36E-03 |
| chemical synaptic transmission (GO:0007268) | 3.80 | 5.43E-06 | 4.16E-03 |
| monoatomic ion transmembrane transport (GO:0034220) | 2.61 | 3.04E-05 | 1.05E-02 |
| inorganic cation transmembrane transport (GO:0098662) | 2.59 | 2.35E-04 | 4.91E-02 |
| organelle organization (GO:0006996) | 0.41 | 5.16E-05 | 1.42E-02 |
| gene expression (GO:0010467) | 0.37 | 2.54E-04 | 4.99E-02 |
| regulation of transcription by RNA polymerase II (GO:0006357) | 0.36 | 5.02E-06 | 4.94E-03 |
| RNA biosynthetic process (GO:0032774) | 0.14 | 1.05E-04 | 2.50E-02 |

## Discussion

Unless they occur as recurrent mutations, DNMs occurring in research participants within a sample will appear as singleton variants and it might seem tempting to use singleton variants to gain information about mutation rates. However in this large cohort each participant carries around 900 singleton variants and is expected to have only 60 or so DNMs, so the vast majority of singleton variants consist of rare variants segregating in the population rather than DNMs. Moreover, as discussed previously, singleton counts can be reduced for variant types with high mutation rates so that overall the counts of singleton variants do not accurately reflect mutation probabilities (Wakeley et al., 2023). Instead, we show that if one considers variants each of which has an allele count of 25, rather than 1, then one finds that the frequencies of the AC25 variant types are very highly correlated with the frequencies of DNMs directly observed in a previously reported, smaller sample (Halldorsson et al., 2019). This finding applies to the present sample of 500,000 genomes and other allele count choices might be more appropriate for other samples.

The large sample size allows for the study of sequence-based effects on relative mutation rates in a normal population. Studying the rates of different mutation types in spermatozoa provides information mainly concerning errors occurring during spermatogenesis (Shoag et al., 2025). Studying cancer cells provides information about somatic mutations which have arisen either during mitosis or at some other point in the cell cycle but which have occurred in the context of disease, when the normal mechanisms for maintaining and accurately copying DNA have been disrupted (Alexandrov et al., 2020). By contrast, AC25 variants observed in a large population sample are expected to reflect mutational events which have occurred in essentially healthy cells. They may have arisen during meiosis or as somatic mutations in primordial germ cells. Thus, studying the relative frequency of different types of AC25 variant may throw light on the molecular mechanisms used by healthy cells to prevent mutations from occurring and this may lead to better insights into susceptibility to neoplastic disease.

We show that the counts of these AC25 variants, which we take to reflect mutation rates, when considered in a trinucleotide context can be very accurately modelled using a small number of mutational signatures which had been obtained from analysing somatic mutations in cancer tissue. The signature with the largest contribution, SBS5, has been described as clock-like, in that the number of these mutations increases with the age of the individual, and may be due to DNA damage to adenine (Alexandrov et al., 2020). It has been reported to dominate among DNMs as well as in somatic cells, cultured cells and tumours and appears to result from multiple types of DNA damage (Spisak et al., 2025). Likewise, the signature with the second largest contribution, SBS1, is also clock-like and may reflect deamination of methylcytosine (Silveira et al., 2024). Comparing these AC25 variant frequencies with those observed in cancer will hopefully facilitate a fuller understanding of pathways to carcinogenesis.

Using variants in the non-transcribed region, we replicate the previously reported finding that the frequency of C>T variants is much increased in the CpG context but we also demonstrate how other aspects of the context can have quite marked effects on variant frequency and hence, we assume, mutation rate (Zhao and Boerwinkle, 2002). Although the trinucleotide context largely predicts mutation rate, we demonstrate that in some situations the fully pentanucleotide context has relevance and we see examples where the more distant bases have two-fold or higher impacts. Investigations into why these specific bases have the effects that they do could lead to a better understanding of the molecular mechanisms involved in generating these mutations.

For some variants, the frequencies differ between non-transcribed and transcribed regions. Possibly this could arise from differences in the organisation of the regions, including features such as repeat content, but these frequency differences might also be the result of mutational processes occurring during transcription. CT[A>G]T* and CA[A>G]T* variants are more frequent in the transcripts of protein coding genes than in the non-transcribed region and occur more frequently on the coding strand than the template strand. The genes with the highest frequencies of CT[A>G]T* variants show enrichment for gene expression and RNA metabolism processes which would be active in all cells, while the genes with the lowest frequencies of these variants show enrichment for processes active only in specialised cells, such as neurons and myocytes. These findings all tend to support the notion that these variants can arise during transcriptional activity and if they occur during embryogenesis or gametocyte differentiation they can then go on to enter the germline.

Some variants differ in frequency between transcripts of protein coding genes and RNA genes. This might reflect differences in genome organisation or transcription levels but such differences might also result from differential effects of the transcription machinery involved in the for the different types of gene.

A more detailed analysis of variant frequencies associated with different types of transcript will be the subject of further work. It will also be fruitful to investigate the distribution of frequencies of variants which have been implicated in human disease. Having a fuller understanding of the mechanisms which can lead to variant formation and how they they are affected by context and location in the genome may assist in improving our ability to predict expected effects and pathogenicity.

## Data availability

The raw data is available on application to UK Biobank at https://ams.ukbiobank.ac.uk/ams/. Any legitimate researcher can apply for access to the UK Biobank dataset by filling in the form at this URL. The counts of variants and of pentanucleotide sequences in the reference genome are provided at https://github.com/davenomiddlenamecurtis/countDNAVariants.

## Acknowledgments

This research has been conducted using the UK Biobank Resource under Application Number 51119. This work uses data provided by patients and collected by NHS England as part of their care and support. This research also used data assets made available by National Safe Haven as part of the Data and Connectivity National Core Study, led by Health Data Research UK in partnership with the Office for National Statistics and funded by UK Research and Innovation (grants MC_PC_20029 and MC_PC_20058). The author wishes to acknowledge the staff supporting the High Performance Computing Cluster, Computer Science Department, University College London. The author wishes to thank the participants who volunteered for the UK Biobank project.

